# A Max-margin Model for Predicting Residue–base Contacts in Protein–RNA Interactions

**DOI:** 10.1101/022459

**Authors:** Shunya Kashiwagi, Kengo Sato, Yasubumi Sakakibara

## Abstract

Protein–RNA interactions (PRIs) are essential for many biological processes, so understanding aspects of the sequences and structures involved in PRIs is important for unraveling such processes. Because of the expensive and time-consuming techniques required for experimental determination of complex protein–RNA structures, various computational methods have been developed to predict PRIs. However, most of these methods focus on predicting only RNA-binding regions in proteins or only protein-binding motifs in RNA. Methods for predicting entire residue–base contacts in PRIs have not yet achieved sufficient accuracy. Furthermore, some of these methods require the identification of 3D structures or homologous sequences, which are not available for all protein and RNA sequences. Here, we propose a prediction method for predicting residue–base contacts between proteins and RNAs using only sequence information and structural information predicted from sequences. The method can be applied to any protein–RNA pair, even when rich information such as its 3D structure, is not available. In this method, residue–base contact prediction is formalized as an integer programming problem. We predict a residue–base contact map that maximizes a scoring function based on sequence-based features such as *k*-mers of sequences and the predicted secondary structure. The scoring function is trained using a max-margin framework from known PRIs with 3D structures. To verify our method, we conducted several computational experiments. The results suggest that our method, which is based on only sequence information, is comparable with RNA-binding residue prediction methods based on known binding data.

## 1. Introduction

Recent studies have begun unraveling the mechanisms of biological processes involving functional non-coding RNAs, most of which interact with RNA-binding proteins (RBPs) in essential roles, such as splicing, transport, localization, and translation. These interactions involve sequence- and structure-specific recognition between proteins and RNAs. Therefore, understanding aspects of sequences and structures involved in protein–RNA interactions (PRIs) is important for understanding many biological processes. To that end, several studies have focused on the analysis and discussion of PRIs [1–3].

Compared with deciphering genomic sequences by using high-throughput sequencing technology, experimental determination of protein–RNA joint structures is both more expensive and more time consuming. Accordingly, rapid computational prediction of PRIs from only sequence information is desirable. Existing methods for computational prediction of PRIs can be roughly classified into four groups. The first group predicts whether a given protein–RNA pair interacts or not [4–7]. A prediction algorithm for this approach can be simply designed from interacting protein–RNA pairs alone, so 3D structures and residue–base contacts are not necessary for use in model training. However, this approach cannot predict binding sites of proteins and RNAs that should be biologically and structurally essential for PRIs. The second group aims to predict RNA-binding residues from protein information. DR_bind1 [8], KYG [9], and OPRA [10] are structure-based methods that use 3D structures from PDB to extract descriptors for prediction. BindN+ [11] and Pprint [12] are sequence-based methods that employ evolutionary information instead of 3D structures. However, this approach ignores the binding partners of target proteins, although some RNA-binding domains in RBPs recognize sequence- and structure-specific motifs in RNA sequences. The third group computes RNA structural motifs recognized by RNA-binding domains in certain proteins and includes MEMERIS [13], RNAcontext [14], CapR [15], and GraphProt [16]. This approach focuses on a certain RBP and extracts RNA motifs as consensus sequences and/or secondary structures of the RBP-binding RNAs. The fourth and final group of methods predicts intermolecular joint structures between proteins and RNAs such as residue–base contacts. To our knowledge, Hayashida *et al*.[17] have developed the only method of this type. However, it is unfortunately not sufficiently accurate.

Accordingly, we propose a prediction method for residue–base contacts between proteins and RNAs based only on sequence information and structural information predicted from sequences. Our method can be applied to any protein–RNA pair, including those for which rich information, such as 3D structures, are unavailable. Residue–base contact prediction is formalized as an integer programming (IP) problem. Our method predicts a residue–base contact map that maximizes a scoring function based on sequence features such as *k*-mers of sequences and predicted secondary structures. The scoring function is trained by a max-margin framework from known PRIs with 3D structures. To verify our method, we performed several computational experiments. The results suggest that our method based on only sequence information is comparable with RNA-binding residue prediction methods based on actual known binding data.

## 2. Methods

We present a novel algorithm for predicting PRIs using IP. Our algorithm consists of the following two parts: (1) prediction of a residue–base contact map given a protein and RNA pair by solving an integer programming problem; and (2) learning a scoring function from a given training dataset using a max-margin framework.

### 2.1. Preliminaries

Let Σ_*p*_ represent the set of 20 canonical amino acid residues and let 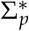 denote the set of all finite amino acid sequences consisting of residues in Σ_*p*_. Similarly, let Σ_*r*_ represent the set of the four canonical ribonucleotide bases (A, C, G, and U) and let 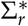 denote the set of all finite RNA sequences consisting of bases in Σ_*r*_. Given a protein 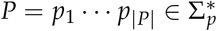 consisting of |*P*| residues and an 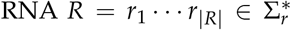 consisting of |*R*| bases, let 𝒞ℳ (*P, R*) represent the space of all possible residue–base contact maps between *P* and *R*. An element *z* ∈ 𝒞ℳ(*P, R*) is represented as an |*P*| *×* |*R*| binary-valued matrix, where *z*_*ij*_ = 1 indicates that residue *p*_*i*_ interacts with the base *r*_*j*_ (Fig. 1). We define the problem of PRI prediction as follows: given a protein *P* and an RNA *R*, predict a residue–base contact map *z* ∈ 𝒞ℳ(*P, R*).

**Figure 1.**
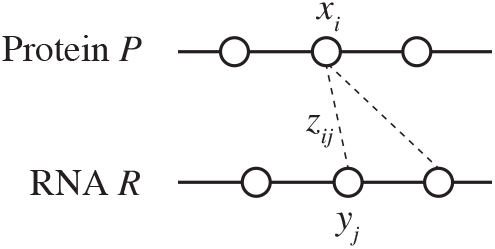
An illustration of binary variables used in the IP formulation.

### 2.2. Scoring model

A scoring model *f* is a function that assigns real-valued scores to protein–RNA pairs (*P, R*) and residue–base contact maps *z* ∈ 𝒞ℳ (*P, R*). Our aim is to find a residue–base contact map *z* ∈ 𝒞ℳ (*P, R*) that maximizes the scoring function *f* (*P, R, z*) for a given protein–RNA pair (*P, R*). The scoring function *f* (*P, R, z*) is computed on the basis of various local features of *P, R*, and *z*. These features correspond to residue features, base features, and residue–base contact features that describe local contexts around residue–base contacts, respectively.

Residue features, as summarized in Table 2, describe the binding preference in the amino acid sequences by local contexts around residue–base contacts. For this purpose, we employ *k*-mers of the amino acids centered on the interacting *i*th residue. For each *k*-mer of the amino acids, 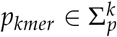, we define a binary-valued local feature of the *i*th residue as

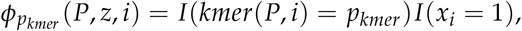

where *I*(*condition*) is an indicator function that takes a value of 1 or 0 depending on whether the *condition* is true or false, *kmer*(*P, i*) is the *k*-mer of the substring of *P* centered on the *i*th residue *p*_*i*_, that is, *kmer*(*P, i*) = *p*_*i* − (*k* − 1)/2_ … *p*_*i*_ … *p*_*i*+(*k* − 1)/2_, and *x*_*i*_ is a binary-valued variable such that *x*_*i*_ = 1 if and only if residue *p*_*i*_ is a binding site (Fig. 1), that is, 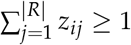. We use *k* = 3 and *k* = 5 to characterize *k*-mer features.

To reduce the sparsity of amino acid contexts, we consider the *k*-mers of simplified alphabets of amino acids proposed by Murphy *et al*. [18], who calculated groups of simplified alphabets based on the BLOSUM50 matrix [19]. Note that Murphy *et al*.[18] have shown that the simplified alphabets are correlated with physiochemical properties such as hydrophobicity, hydrophilicity, and polarity, which may have important roles in PRIs. We employ the simplified alphabets of 10 groups, Σ_*g*10_, and those of 4 groups, Σ_*g*4_ (Table 1).

**Table 1.** Groups of amino acids as defined by Murphy *et al*. [18]

|  | # | groups |
| --- | --- | --- |
| $\Sigma_{g10}$ | 10 | LVIM, C, A, G, ST, P, FYW, EDNQ, KR, H |
| $\Sigma_{g4}$ | 4 | LVIMC, AGSTP, FYW, EDNQKRH |

**Table 2.** A summary of residue features

| Type | Context len. | # of features |
| --- | --- | --- |
| Residues | 3 | $20^3$ |
| | 5 | $20^5$ |
| Simplified alphabets (10 groups) | 5 | $10^5$ |
| | 7 | $10^7$ |
| Simplified alphabets (4 groups) | 5 | $4^5$ |
| | 7 | $4^7$ |
| Secondary structures | 3 | $8^3$ |
| | 5 | $8^5$ |

For each string 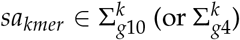, we define a binary-valued local feature of the *i*th residue as

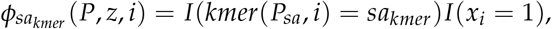

where *P*_*sa*_ is the string of simplified alphabets Σ_*g*10_ (or Σ_*g*4_) converted from *P* according to Table 1. In contrast with the *k*-mers used in other part of this algorithm, we instead use *k* = 5 and *k* = 7 for the *k*-mers of simplified alphabets.

To consider the structural preference of RNA-binding residues, we employ secondary structures predicted by SSpro8 [20]. We predict one structural element [*α*-helix (H), 3-helix (G), 5-helix (I), folded (E), *β*-turn (B), corner (T), curl (S), and loop (–)] for each residue. For each string *sp*_*kmer*_ of structural elements of length *k*, we define a binary-valued local feature of the *i*th residue as

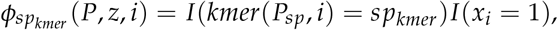

where *P*_*sp*_ is the string of structural elements predicted from *P*. Here, we again use structural contexts with lengths *k* = 3 and *k* = 5.

The collection of occurrences of the residue features are calculated as

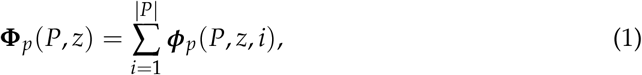

where ***ϕ***_*p*_(*P, z, i*) is a vector whose elements are the residue features of the *i*th residue mentioned above.

Base features, as summarized in Table 3, describe the binding preference in the ribonucleotide sequences by local contexts around residue–base contacts. In addition to the residue features, we employ the *k*-mer contexts of the ribonucleotides centered on the interacting *j*th base. For each *k*-mer of the ribonucleotides 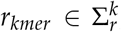, we define a binary-valued local feature of the *j*th base as

**Table 3.** A summary of base features

| Type | Context len. | # of features |
| --- | --- | --- |
| Bases | 3 | $4^3$ |
| | 5 | $4^5$ |
| Secondary structures | 3 | $6^3$ |
| | 5 | $6^5$ |

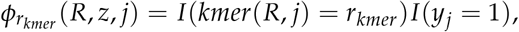

where *y*_*j*_ is a binary-valued variable such that *y*_*j*_ = 1 if and only if the residue *r*_*j*_ is a binding site (Fig. 1), that is, 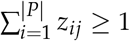. Here, we once again use *k* = 3 and 5 for the *k*-mer features.

To consider the structural preference of binding sites, we employ secondary structures predicted by CentroidFold [21]. We assign a structural element [external loop (E), hairpin loop (H), internal loop (I), bulge (B), multibranch loop (M), or stack (S), as shown in Fig. 2] to each base. Note that to encode secondary structures as a sequence, this encoding of structural profiles loses a portion of the structural information, e.g., basepairing partners for stacking bases. However, this approach is still efficient for describing structural information [13–15]. For each *k*-length string *sr*_*kmer*_ of structural elements, we define a binary-valued local feature of the *j*th base as

**Figure 2.**
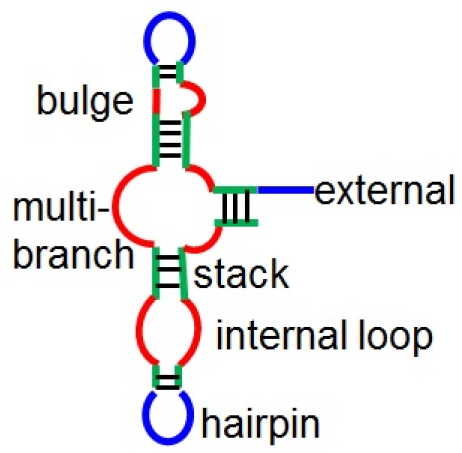
Structural elements in RNA secondary structures.

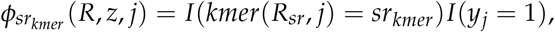

where *R*_*sr*_ is the string of structural elements predicted from *R*. Here, we use structural contexts with lengths *k* = 3 and *k* = 5.

The collection of occurrences of the base features are calculated as

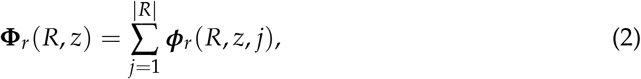

where ***ϕ***_*r*_(*R, z, j*) is a vector whose elements are the base features of the *j*th base mentioned above.

Residue–base contact features, which are summarized in Table 4, describe the binding affinity between the local contexts of amino acids and ribonucleotides. For this purpose, we employ combinations of the residue features and the base features mentioned above. For example, for each pair of *k*-mers of amino acids *p*_*kmer*_ and ribonucleotides *r*_*kmer*_, we define a binary-valued local feature of the *i*th residue and the *j*th base:

**Table 4.**
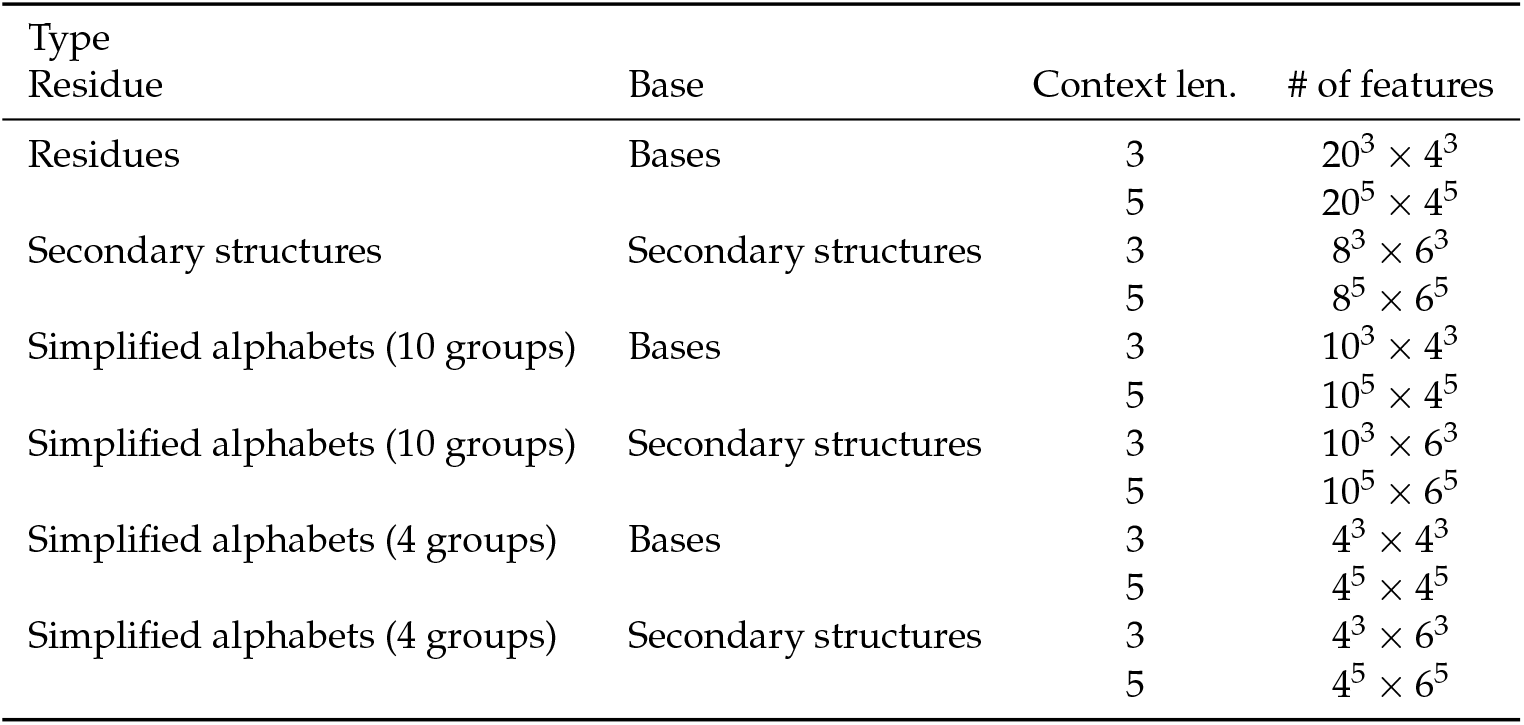
A summary of residue–base contact features

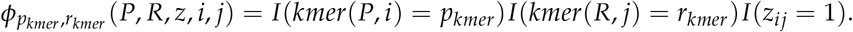

The collection of occurrences of the residue–base contact features are calculated as

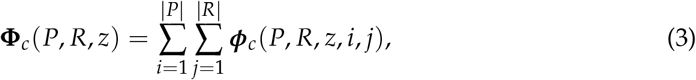

where ***ϕ***_*c*_(*P, R, z, i, j*) is a vector whose elements are the residue–base contact features of the *i*th residue and the *j*th base mentioned above.

The notation **Φ**(*P, R, z*) denotes the feature representation of protein–RNA pair (*P, R*) and its residue–base contact map *z* ∈ 𝒞ℳ (*P, R*), that is, the collection of occurrences of local features in *P, R*, and *z* defined as follows:

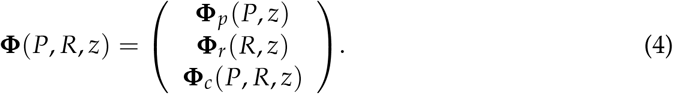

Each feature in **Φ** is associated with a corresponding parameter, and the score for the feature is defined as the value of the occurrence multiplied by the corresponding parameter. We define the scoring model *f* (*P, R, z*) as a linear function

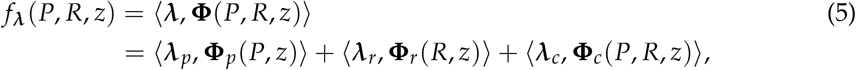

where ⟨·,·⟩ is the inner product and 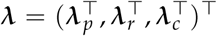 is the corresponding parameter vector trained with training data as described in Sec. 2.4.

### 2.3. IP formulation

To formulate the problem as an IP problem, we rewrite the scoring function (5) as

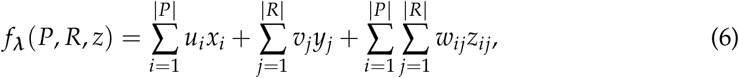

where *u*_*i*_, *v*_*i*_, and *w*_*ij*_ represent the binding preferences for *x*_*i*_, *y*_*j*_, and *z*_*ij*_, respectively, calculated as

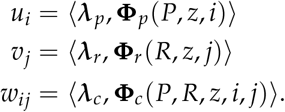

We find a *z* ∈ 𝒞ℳ (*P, R*) that maximizes the objective function (6) under the following constraints to ensure consistency among the variables *x*_*i*_, *y*_*j*_, and *z*_*ij*_ as follows:

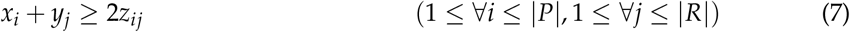

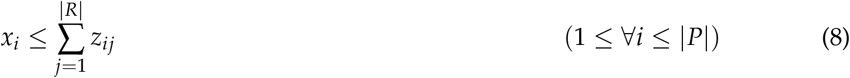

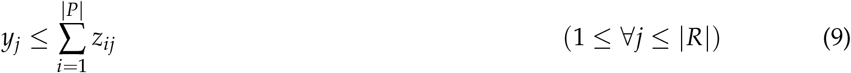

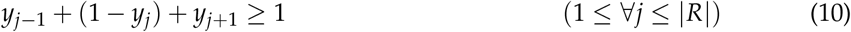

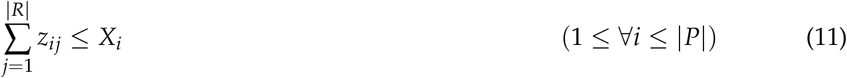

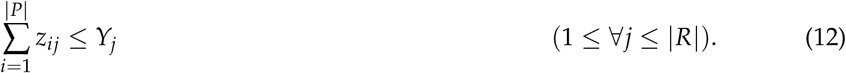

The constraints defined by Eqs. (7)–(9) describe the relation between contacts *z*_*ij*_ and binding sites *x*_*i*_, *y*_*j*_. The constraint defined by Eq. (10) disallows any isolated interacting bases, which are rare in PRIs. The constraints defined by Eqs. (11) and (12) define the upper bound on the number of contacts *X*_*i*_ and *Y*_*j*_ for each residue and base, respectively.

### 2.4. Learning algorithm

To optimize feature parameter ***λ***, we employ a max-margin framework called structured support vector machines [22]. Given a training dataset 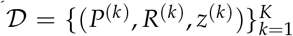, where *P*^(*k*)^ and *R*^(*k*)^ are the protein and RNA sequences, respectively, and *z*^(*k*)^ ∈ 𝒞ℳ (*P*^(*k*)^, *R*^(*k*)^) is their corresponding contact map for the *k*th datapoint, we aim to find the parameter ***λ*** that minimizes the objective function

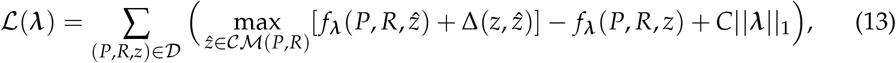

where ||.||_1_ is the *ℓ*_1_ norm and *C* is a weight for the *ℓ*_1_ regularization term to avoid overfitting to the training data. Here, 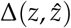 is a loss function of 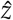 for *z* defined as

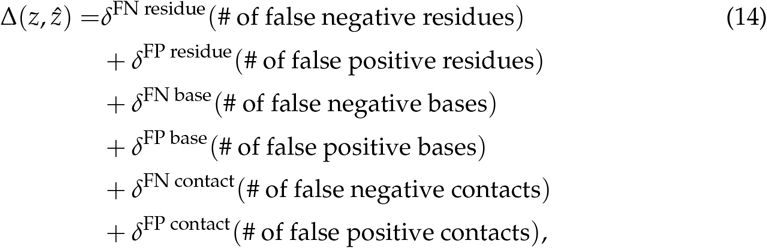

where *δ*^FN residue^, *δ*^FP residue^, *δ*^FN base^, *δ*^FP base^, *δ*^FN contact^, and *δ*^FP contact^ are hyperparameters controlling the trade-off between sensitivity and specificity for learning the parameters. In this case, we can calculate the first term of Eq. (13) by replacing scores *u*_*i*_, *v*_*j*_, and *w*_*ij*_ in Eq. (6) as follows:

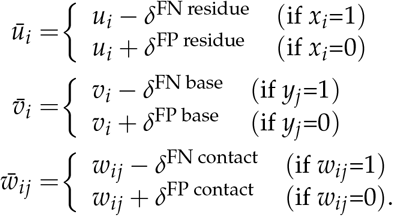

See Sec. S1 in the Supplementary Material for the derivation.

To minimize the objective function (13), we can apply stochastic subgradient descent (Fig. 3) or forward-backward splitting [23].

**Figure 3.**
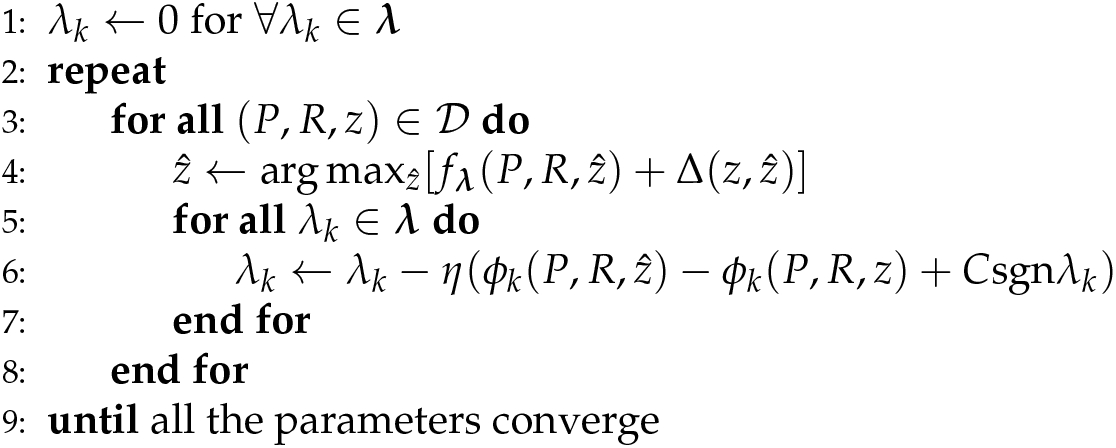
The stochastic subgradient descent algorithm for a structured support vector machine; sgn is the sign function, whereas *η* > 0 is the predefined learning rate.

## 3. Results

### 3.1. Implementation

Our method was implemented using the IBM CPLEX optimizer^1^ for solving IP problems (6)–(12). To extract the structural feature elements described in Sec. 2.2, we employed SSpro8 [20] and CentroidFold [21] to predict secondary structures of protein and RNA sequences, respectively. We empirically chose the following hyperparameters: penalty for positives, *δ*^FN *^ = 0.5; penalty for negatives, *δ*^FP *^ = 0.005; and the weight for the *ℓ*_1_ regularization term, *C* = 10^−5^. See Sec. S2 in the Supplementary Material for details. We implemented AdaGrad [24] to control the learning rate *η* in Fig. 3. The source code for our algorithm is available at https://github.com/keio-bioinformatics/practip.

### 3.2. Dataset

We prepared our datasets in accordance with those of Chen *et al*. [8] and Miao *et a*. [25] and extracted RNA-bound proteins with an X-ray resolution of ≤ 3.0 Å from the Protein Data Bank (PDB) [26]. To reduce dataset redundancy, we discarded some extracted data such that the dataset contained no protein pairs whose sequence identity was > 30%. As a result, our test dataset consisted of 98 protein–RNA interacting pairs from 81 protein–RNA complexes from Chen *et al*. [8], and our training dataset consisted of 4399 protein–RNA interacting pairs from 772 protein–RNA complexes was from Miao *et al*. [25]. We considered a residue to bind RNA if at least one non-hydrogen atom was contained within the van der Waals contact (4.0Å) or hydrogen-bonding distance (3.5Å) from the non-hydrogen atom of its binding partner. We employed HBPLUS [27] to detect the hydrogen bonds and van der Waals contacts.

### 3.3. Prediction of residue–base contacts

To validate our method, we conducted computational experiments on our dataset, comparing the accuracy under several conditions related to the maximum number of contacts for each residue and base, *X*_*i*_ and *Y*_*j*_ in Eqs. (11) and (12) from 1 to 9, and no upper bounds.

We evaluated the accuracy of predicting residue–base contacts between proteins and RNAs using three measures: predicted residue–base contacts, binding residues in proteins, and binding bases in RNA sequences. The accuracy of residue–base contacts is assessed by the positive predictive value (PPV) and the sensitivity (SEN), respectively defined as

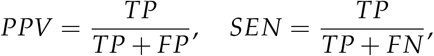

where *TP* is the number of correctly predicted contacts (true positives), *FP* is the number of incorrectly predicted contacts (false positives), and *FN* is the number of contacts in the true contact map that were not predicted (false negatives). We also used the F-value as a balanced measure between PPV and SEN, and it is defined as their harmonic mean:

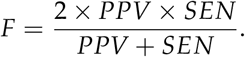

The accuracy of binding residues and binding bases is defined in the same way.

Table 5 shows the accuracy of predicting residue–base contacts in PRIs, binding residues in proteins, and binding bases in RNA sequences for upper bounds of contacts *X*_*i*_, *ϒ*_*j*_ in Eqs. (11) and (12) from 1 to 9 and for no upper bounds. The case with the strongest constraint (*X*_*i*_ = *ϒ*_*j*_ = 1) has a very high PPV because it limits the number of contacts to be predicted, while its SEN is poor because of a lack of coverage of the prediction. On the other hand, if there is no constraint on the number of contacts (corresponding to the row labeled “no limit” in Table 5), both PPV and SEN are not high owing to many incorrect predictions being made. We found that if the upper limit of the number of contacts is set between 4 and 9, reasonably accurate contact prediction, residue binding site prediction, and base binding site prediction can be obtained. As a result, we set *X*_*i*_ = *ϒ*_*j*_ = 8 as the default constraint for the upper bound of the number of contacts.

**Table 5.** Accuracy under varying conditions on the maximum number of contacts for each residue and base.

| upper bounds<br>of # contacts ($X_i, Y_j$) | Contacts | | | Binding residues | | | Binding bases | | |
| --- | --- | --- | --- | --- | --- | --- | --- | --- | --- |
|  | PPV | SEN | F | PPV | SEN | F | PPV | SEN | F |
| 1 | **0.599** | 0.192 | 0.278 | **0.829** | 0.349 | 0.460 | **0.877** | 0.361 | 0.481 |
| 2 | 0.552 | 0.347 | 0.414 | 0.736 | 0.509 | 0.578 | 0.796 | 0.506 | 0.597 |
| 3 | 0.523 | 0.436 | 0.462 | 0.679 | 0.585 | 0.608 | 0.744 | 0.595 | 0.644 |
| 4 | 0.532 | 0.480 | 0.491 | 0.676 | 0.642 | 0.638 | 0.718 | 0.626 | 0.656 |
| 5 | 0.534 | 0.506 | 0.507 | 0.655 | 0.667 | 0.641 | 0.693 | **0.656** | **0.657** |
| 6 | 0.537 | 0.515 | 0.514 | 0.669 | 0.671 | 0.654 | 0.688 | 0.647 | 0.652 |
| 7 | 0.541 | 0.520 | 0.518 | 0.671 | 0.685 | 0.663 | 0.677 | 0.649 | 0.647 |
| 8 | 0.539 | **0.525** | **0.519** | 0.664 | **0.688** | **0.657** | 0.684 | 0.655 | 0.652 |
| 9 | 0.531 | 0.513 | 0.510 | 0.658 | 0.679 | 0.649 | 0.659 | 0.650 | 0.638 |
| no limit | 0.321 | 0.367 | 0.328 | 0.481 | 0.556 | 0.493 | 0.535 | 0.530 | 0.508 |

It should be noted that in this experiment, we were unable to compare our method with the method by Hayashida *et al*. [17], which is the only published method for predicting reside–base contacts in PRIs. Specifically, we were unable to conduct an experiment using the method by Hayashida *et al*. on the same dataset because their software implementation is not yet available and their method requires homologous sequences with accurate alignments to calculate evolutionary information. In addition, Hayashida *et al*. [17] have reported that the method is not sufficiently accurate for such analyses.

### 3.4. Comparison of binding residues predictions among the present and existing methods

We compared our method with existing methods for predicting RNA-binding residues in proteins. DR_bind1 [8], KYG [9], and OPRA [10] are structure-based methods that use 3D structures from PDB to extract descriptors for prediction. BindN+ [11] and Pprint [12] are sequence-based methods that employ evolutionary information instead of 3D structures. Table 6 indicates that our method is comparable to other methods. Recall that our method employs only sequence information and structural information predicted from sequences as well as information on the partner RNAs bound to RNA-binding proteins, rather than 3D structures and evolutionary information.

**Table 6.**
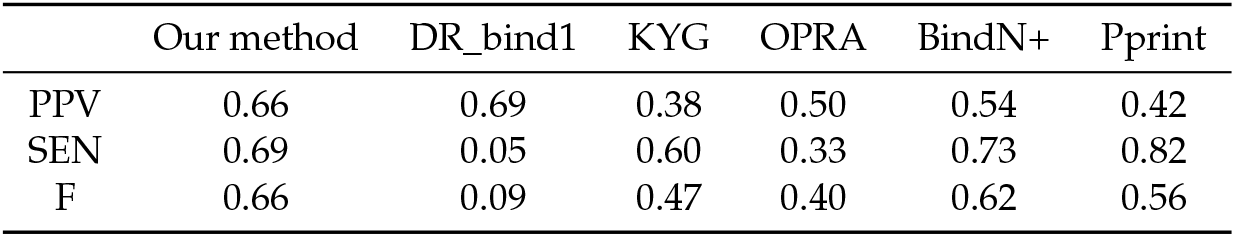
Comparison of our method with other existing methods on our dataset.

|  | Our method | DR_bind1 | KYG | OPRA | BindN+ | Pprint |
| --- | --- | --- | --- | --- | --- | --- |
| PPV | 0.66 | 0.69 | 0.38 | 0.50 | 0.54 | 0.42 |
| SEN | 0.69 | 0.05 | 0.60 | 0.33 | 0.73 | 0.82 |
| F | 0.66 | 0.09 | 0.47 | 0.40 | 0.62 | 0.56 |

## 4. Discussion

Several existing methods for predicting PRIs utilize evolutionary information from homologous sequences, [11,12] for protein sequences and [17] for both protein and RNA sequences. Homologous sequences of target sequences are typically searched for in large databases using a highly sensitive homology search engine such as PSI-BLAST [28]. Furthermore, to extract evolutionary information, homologous sequences must be aligned before PRI prediction. Homology searches are employed in a wide range of analyses, such as functional analysis of proteins, because if homologous proteins can be found in curated databases, the function of the target protein can be easily inferred. However, as described above and by Zhang *et al*. [29], the secondary structures of proteins play essential roles in residue–base contacts. Similarly, structural elements of RNA secondary structures also serve as key descriptors for residue–base contact prediction [13–16]. This means that structure-based homology searches are needed for PRI prediction based on evolutionary information. Although efficient structural alignment algorithms for proteins (e.g., [30]) and RNAs (e.g., [31]) have recently been developed, they have not yet been successfully applied to large-scale homology searches.

To our knowledge, Hayashida *et al*. [17] have developed the only existing method that predicts intermolecular joint structures between proteins and RNAs such as residue–base contacts; however, this method is unfortunately not sufficiently accurate. The method by Hayashida *et al*. [17] is similar to our method in that its approach is based on a machine learning technique with *ℓ*_1_ regularization. The main difference between our method and the method by Hayashida *et al*. [17] is that our method employs a large number of features, including structural information about proteins and RNAs, which have been shown to serve as key descriptors of PRIs as mentioned above.

We utilized the structural profiles of predicted RNA secondary structures, which does lose an important part of structural information, such as base-pairing partners for stacking bases. Most of the existing RBP-binding RNA motif finding methods [13–15] have also utilized similar encoding, which may not be suitable for dealing with the recognition sites of double-stranded RNA-binding proteins. GraphProt [16] is an exceptional algorithm that utilizes graph-based encoding of RNA secondary structures. Our method should be extended by utilizing another structural profile with no loss of base pairing information like the graph-based encoding of GraphProt.

As shown in Sec. 2.3, we formulated the residue–base contact prediction as an IP problem, which enables us to build a flexible model, including, for example, constraints on the upper bound on the number of contacts for each residue and base. In contrast to the RNA–RNA interaction model in which each base interacts with at most one base via hydrogen bonds such as Watson–Crick and wobble base pairs, PRIs contain diverse patterns of residue–base contacts. For example, Kondo *et al*. have classified residue–base contacts with respect to three interaction edges on nucleotides (Watson–Crick, Hoogsteen, and sugar) with side-chains and backbones of their partner residues, and have analyzed their propensities [1]. Thus, there is room for further improvement of our model, which can be extended by using other constraints for each contact between a residue and a base to include such considerations.

The large-scale sequencing data produced by RNA-related high-throughput sequencing technologies, such as Structure-seq [32] and hiCLIP [33], will help us improve our algorithm, especially by providing data for training the model. In the present work, we employed complete joint 3D structures of proteins and RNAs as the training dataset, which was not sufficiently large. We cannot build from large-scale sequencing data a complete dataset with residue–base contact maps, but we can partially calculate structural profiles and binding bases from *in vivo* chemical probing data such as Structure-seq datasets. This information will significantly help us improve our model.

Deep learning has been increasingly used in various fields, including bioinformatics, in recent years. Wei *et al*. [34] have provided a review of the use of deep learning in RNA– protein interaction prediction. Yamada *et al*. [35] have developed a method to accurately identify RNA sequences that interact with a particular protein by using the DNABERT model [36] that is pre-trained using the human genome. Although our method does not use deep learning, we expect to achieve higher accuracy in prediction by using a pre-trained BERT model, which could be improved through the application of deep learning relatively easily.

## 5. Conclusion

We developed a max-margin framework for predicting residue–base contacts between proteins and RNAs based on integer programming. To verify our method, we performed several computational experiments. The results suggest that our method based only on sequence information and structural information predicted from sequences is comparable with RNA-binding residue prediction methods based on known binding data. Further improvements are needed, such as the incorporation of informative features, the development of a joint prediction model that simultaneously predicts RNA secondary structures and protein contact maps, and the utilization of high-throughput sequencing data that can deal with PRI without residue–base contact information as training data.

## Author Contributions

Conceptualization, K.S.; methodology, K.S.; software, S.K. and K.S.; resources, S.K. and K.S.; data curation, S.K.; writing—original draft preparation, S.K. and K.S.; writing—review and editing, K.S. and Y.S.; supervision, K.S. and Y.S.; project administration, K.S.; funding acquisition, K.S. and Y.S. All authors have read and agreed to the publication of this version of the manuscript.

## Acknowledgments

The supercomputer system used for this research was made available by the National Institute of Genetics (NIG), Research Organization of Information and Systems (ROIS).

## Abbreviations

The following abbreviations are used in this manuscript:

IP: Integer programming
PPV: Positive predictive value
PRI: Protein–RNA interaction
RBP: RNA binding protein
SEN: Sensitivity
SP: Structural profile
SVM: Support vector machine

## Footnotes

1 http://www.ibm.com/software/integration/optimization/cplex-optimizer/

## References

1. Kondo, J.; Westhof, E. Classification of pseudo pairs between nucleotide bases and amino acids by analysis of nucleotide-protein complexes. Nucleic Acids Res. 2011, 39, 8628–8637.

2. Iwakiri, J.; Tateishi, H.; Chakraborty, A.; Patil, P.; Kenmochi, N. Dissecting the protein-RNA interface: the role of protein surface shapes and RNA secondary structures in protein-RNA recognition. Nucleic Acids Res. 2012, 40, 3299–3306.

3. Iwakiri, J.; Kameda, T.; Asai, K.; Hamada, M. Analysis of base-pairing probabilities of RNA molecules involved in protein-RNA interactions. Bioinformatics 2013, 29, 2524–2528.

4. Pancaldi, V.; Bahler, J. In silico characterization and prediction of global protein-mRNA interactions in yeast. Nucleic Acids Res. 2011, 39, 5826–5836.

5. Muppirala, U.K.; Honavar, V.G.; Dobbs, D. Predicting RNA-protein interactions using only sequence information. BMC Bioinformatics 2011, 12, 489.

6. Bellucci, M.; Agostini, F.; Masin, M.; Tartaglia, G.G. Predicting protein associations with long noncoding RNAs. Nat. Methods 2011, 8, 444–445.

7. Wang, Y.; Chen, X.; Liu, Z.P.; Huang, Q.; Wang, Y.; Xu, D.; Zhang, X.S.; Chen, R.; Chen, L. De novo prediction of RNA-protein interactions from sequence information. Mol Biosyst 2013, 9, 133–142.

8. Chen, Y.C.; Sargsyan, K.; Wright, J.D.; Huang, Y.S.; Lim, C. Identifying RNA-binding residues based on evolutionary conserved structural and energetic features. Nucleic Acids Res. 2014, 42, e15.

9. Kim, O.T.; Yura, K.; Go, N. Amino acid residue doublet propensity in the protein-RNA interface and its application to RNA interface prediction. Nucleic Acids Res. 2006, 34, 6450–6460.

10. Perez-Cano, L.; Fernandez-Recio, J. Optimal protein-RNA area, OPRA: a propensity-based method to identify RNA-binding sites on proteins. Proteins 2010, 78, 25–35.

11. Wang, L.; Huang, C.; Yang, M.Q.; Yang, J.Y. BindN+ for accurate prediction of DNA and RNA-binding residues from protein sequence features. BMC Syst Biol 2010, 4 Suppl 1, S3.

12. Kumar, M.; Gromiha, M.M.; Raghava, G.P. Prediction of RNA binding sites in a protein using SVM and PSSM profile. Proteins 2008, 71, 189–194.

13. Hiller, M.; Pudimat, R.; Busch, A.; Backofen, R. Using RNA secondary structures to guide sequence motif finding towards single-stranded regions. Nucleic Acids Res. 2006, 34, e117.

14. Kazan, H.; Ray, D.; Chan, E.T.; Hughes, T.R.; Morris, Q. RNAcontext: a new method for learning the sequence and structure binding preferences of RNA-binding proteins. PLoS Comput. Biol. 2010, 6, e1000832.

15. Fukunaga, T.; Ozaki, H.; Terai, G.; Asai, K.; Iwasaki, W.; Kiryu, H. CapR: revealing structural specificities of RNA-binding protein target recognition using CLIP-seq data. Genome Biol. 2014, 15, R16.

16. Maticzka, D.; Lange, S.J.; Costa, F.; Backofen, R. GraphProt: modeling binding preferences of RNA-binding proteins. Genome Biol. 2014, 15, R17.

17. Hayashida, M.; Kamada, M.; Song, J.; Akutsu, T. Prediction of protein-RNA residue-base contacts using two-dimensional conditional random field with the lasso. BMC Syst Biol 2013, 7 Suppl 2, S15.

18. Murphy, L.R.; Wallqvist, A.; Levy, R.M. Simplified amino acid alphabets for protein fold recognition and implications for folding. Protein Eng. 2000, 13, 149–152.

19. Henikoff, S.; Henikoff, J.G. Amino acid substitution matrices from protein blocks. Proc. Natl. Acad. Sci. U.S.A. 1992, 89, 10915–10919.

20. Magnan, C.N.; Baldi, P. SSpro/ACCpro 5: almost perfect prediction of protein secondary structure and relative solvent accessibility using profiles, machine learning and structural similarity. Bioinformatics 2014, 30, 2592–2597. doi:10.1093/bioinformatics/btu352.

21. Hamada, M.; Kiryu, H.; Sato, K.; Mituyama, T.; Asai, K. Prediction of RNA secondary structure using generalized centroid estimators. Bioinformatics 2009, 25, 465–473.

22. Tsochantaridis, I.; Joachims, T.; Hofmann, T.; Altun, Y. Large Margin Methods for Structured and Interdependent Output Variables. J. Mach. Learn. Res. 2005, 6, 1453–1484.

23. Duchi, J.; Singer, Y. Efficient online and batch learning using forward backward splitting. Journal of Machine Learning Research 2009, 10, 2899–2934.

24. Duchi, J.; Hazan, E.; Singer, Y. Adaptive subgradient methods for online learning and stochastic optimization. The Journal of Machine Learning Research 2011, 12, 2121–2159.

25. Miao, Z.; Westhof, E. A Large-Scale Assessment of Nucleic Acids Binding Site Prediction Programs. PLoS Comput. Biol. 2015, 11, e1004639. doi:10.1371/journal.pcbi.1004639.

26. Rose, P.W.; Beran, B.; Bi, C.; Bluhm, W.F.; Dimitropoulos, D.; Goodsell, D.S.; Prlic, A.; Quesada, M.; Quinn, G.B.; Westbrook, J.D.; Young, J.; Yukich, B.; Zardecki, C.; Berman, H.M.; Bourne, P.E. The RCSB Protein Data Bank: redesigned web site and web services. Nucleic Acids Res. 2011, 39, 392–401.

27. McDonald, I.K.; Thornton, J.M. Satisfying hydrogen bonding potential in proteins. J. Mol. Biol. 1994, 238, 777–793.

28. Altschul, S.F.; Madden, T.L.; Schaffer, A.A.; Zhang, J.; Zhang, Z.; Miller, W.; Lipman, D.J. Gapped BLAST and PSI-BLAST: a new generation of protein database search programs. Nucleic Acids Res. 1997, 25, 3389–3402.

29. Zhang, T.; Zhang, H.; Chen, K.; Ruan, J.; Shen, S.; Kurgan, L. Analysis and prediction of RNA-binding residues using sequence, evolutionary conservation, and predicted secondary structure and solvent accessibility. Curr. Protein Pept. Sci. 2010, 11, 609–628.

30. Deng, X.; Cheng, J. MSACompro: protein multiple sequence alignment using predicted secondary structure, solvent accessibility, and residue-residue contacts. BMC Bioinformatics 2011, 12, 472.

31. Sato, K.; Kato, Y.; Akutsu, T.; Asai, K.; Sakakibara, Y. DAFS: simultaneous aligning and folding of RNA sequences via dual decomposition. Bioinformatics 2012, 28, 3218–3224.

32. Ding, Y.; Tang, Y.; Kwok, C.K.; Zhang, Y.; Bevilacqua, P.C.; Assmann, S.M. In vivo genome-wide profiling of RNA secondary structure reveals novel regulatory features. Nature 2014, 505, 696–700.

33. Sugimoto, Y.; Vigilante, A.; Darbo, E.; Zirra, A.; Militti, C.; D’Ambrogio, A.; Luscombe, N.M.; Ule, J. hiCLIP reveals the in vivo atlas of mRNA secondary structures recognized by Staufen 1. Nature 2015, 519, 491–494.

34. Wei, J.; Chen, S.; Zong, L.; Gao, X.; Li, Y. Protein-RNA interaction prediction with deep learning: Structure matters 2021. [q-bio.BM/2107.12243].

35. Yamada, K.; Hamada, M. Prediction of RNA-protein interactions using a nucleotide language model. doi:10.1101/2021.04.27.441365.

36. Ji, Y.; Zhou, Z.; Liu, H.; Davuluri, R.V. DNABERT: pre-trained Bidirectional Encoder Representations from Transformers model for DNA-language in genome. Bioinformatics 2021. doi:10.1093/bioinformatics/btab083.

