## Supplemental Information for "A Max-margin Model for Predicting Residue–base Contacts in Protein–RNA Interactions"

#### S1 Derivation of scoring functions for max-margin training

The loss function of the prediction $\hat{z}$ against the positive data $z$ (Eq. (14) in the main paper) can be transformed into the following using binary-valued variables:

$$
\begin{aligned}
\Delta(z,\hat{z}) =& \delta^{\text{FN residue}}(\text{\# of false negative residues})\\
&+\delta^{\text{FP residue}}(\text{\# of false positive residues})\\
&+\delta^{\text{FN base}}(\text{\# of false negative bases})\\
&+\delta^{\text{FP base}}(\text{\# of false positive bases})\\
&+\delta^{\text{FN contact}}(\text{\# of false negative contacts})\\
&+\delta^{\text{FP contact}}(\text{\# of false positive contacts})\\
=& \delta^{\text{FN residue}}\sum_{i=1}^{N_p} I(x_i=1)I(\hat{x}_i=0) + \delta^{\text{FP residue}}\sum_{i=1}^{N_p} I(x_i=0)I(\hat{x}_i=1)\\
&+\delta^{\text{FN base}}\sum_{j=1}^{N_r} I(y_j=1)I(\hat{y}_j=0) + \delta^{\text{FP base}}\sum_{j=1}^{N_r} I(y_j=0)I(\hat{y}_j=1)\\
&+\delta^{\text{FN contact}}\sum_{i=1}^{N_p}\sum_{j=1}^{N_r} I(z_{ij}=1)I(\hat{z}_{ij}=0) + \delta^{\text{FP contact}}\sum_{i=1}^{N_p}\sum_{j=1}^{N_r} I(z_{ij}=0)I(\hat{z}_{ij}=1)\\
=& \sum_{i=1}^{N_p}\left\{\delta^{\text{FN residue}}x_i(1-\hat{x}_i) + \delta^{\text{FP residue}}(1-x_i)\hat{x}_i\right\}\\
&+\sum_{j=1}^{N_r}\left\{\delta^{\text{FN base}}y_j(1-\hat{y}_j) + \delta^{\text{FP base}}(1-y_j)\hat{y}_j\right\}\\
&+\sum_{i=1}^{N_p}\sum_{j=1}^{N_r}\left\{\delta^{\text{FN contact}}z_{ij}(1-\hat{z}_{ij}) + \delta^{\text{FP contact}}(1-z_{ij})\hat{z}_{ij}\right\}.
\end{aligned}
$$

Here, $I(x_i=1)I(\hat{x}_i=0)=1$ if $\hat{x}_i$ is a false negative and 0 otherwise, and $I(x_i=0)I(\hat{x}_i=1)=1$ if $\hat{x}_i$ is a false positive and 0 otherwise. We also use the fact that $I(x_i=1)=x_i$ and $I(x_i=0)=1-x_i$.

Therefore, the first term of Eq. (13) in the main paper can be simplified into:

$$
\begin{aligned}
f_\lambda(P,R,\hat{z}) + \Delta(z,\hat{z}) = & \sum_{i=1}^{N_p} u_i \hat{x}_i + \sum_{j=1}^{N_r} v_j \hat{y}_j + \sum_{i=1}^{N_p}\sum_{j=1}^{N_r} w_{ij}\hat{z}_{ij} \\
& + \sum_{i=1}^{N_p} \left\{ \delta^{\text{FN residue}} x_i (1-\hat{x}_i) + \delta^{\text{FP residue}} (1-x_i)\hat{x}_i \right\} \\
& + \sum_{j=1}^{N_r} \left\{ \delta^{\text{FN base}} y_j (1-\hat{y}_j) + \delta^{\text{FP base}} (1-y_j)\hat{y}_j \right\} \\
& + \sum_{i=1}^{N_p}\sum_{j=1}^{N_r} \left\{ \delta^{\text{FN contact}} z_{ij} (1-\hat{z}_{ij}) + \delta^{\text{FP contact}} (1-z_{ij})\hat{z}_{ij} \right\} \\
= & \sum_{i=1}^{N_p} \left\{ \left[ u_i - \delta^{\text{FN residue}} x_i + \delta^{\text{FP residue}} (1-x_i) \right] \hat{x}_i + \delta^{\text{FN residue}} x_i \right\} \\
& + \sum_{j=1}^{N_r} \left\{ \left[ v_j - \delta^{\text{FN base}} y_j + \delta^{\text{FP base}} (1-y_j) \right] \hat{y}_j + \delta^{\text{FN base}} y_j \right\} \\
& + \sum_{i=1}^{N_p}\sum_{j=1}^{N_r} \left\{ \left[ w_{ij} - \delta^{\text{FN contact}} z_{ij} + \delta^{\text{FP contact}} (1-z_{ij}) \right] \hat{z}_{ij} + \delta^{\text{FN contact}} z_{ij} \right\} \\
= & \sum_{i=1}^{N_p} \bar{u}_i \hat{x}_i + \sum_{j=1}^{N_r} \bar{v}_j \hat{y}_j + \sum_{i=1}^{N_p}\sum_{j=1}^{N_r} \bar{w}_{ij}\hat{z}_{ij} + Const,
\end{aligned}
$$

where

$$
\begin{aligned}
\bar{u}_i = & u_i - \delta^{\text{FN residue}} x_i + \delta^{\text{FP residue}} (1-x_i) \\
= & \begin{cases} u_i - \delta^{\text{FN residue}} & (\text{if } x_i{=}1) \\ u_i + \delta^{\text{FP residue}} & (\text{if } x_i{=}0) \end{cases} \\
\bar{v}_i = & v_j - \delta^{\text{FN base}} y_j + \delta^{\text{FP base}} (1-y_j) \\
= & \begin{cases} v_i - \delta^{\text{FN base}} & (\text{if } y_j{=}1) \\ v_i + \delta^{\text{FP base}} & (\text{if } y_j{=}0) \end{cases} \\
\bar{w}_{ij} = & w_{ij} - \delta^{\text{FN contact}} z_{ij} + \delta^{\text{FP contact}} (1-z_{ij}) \\
= & \begin{cases} w_{ij} - \delta^{\text{FN contact}} & (\text{if } z_{ij}{=}1) \\ w_{ij} + \delta^{\text{FP contact}} & (\text{if } z_{ij}{=}0) \end{cases} \\
Const = & \sum_{i=1}^{N_p} \delta^{\text{FN residue}} x_i + \sum_{j=1}^{N_r} \delta^{\text{FN base}} y_j + \sum_{i=1}^{N_p}\sum_{j=1}^{N_r} \delta^{\text{FN contact}} z_{ij}
\end{aligned}
$$

The last equation indicates that we can maximize the first term of the objective function (13) by replacing scores $u_i, v_j$ and $w_{ij}$ with a constant difference $Const$ that is independent of $\hat{x}_i, \hat{y}_j$ and $\hat{z}_{ij}$.

#### S2 Hyperparameters

We empirically chose the hyperparameters for the max-margin training: the penalty for positives $\delta^{\text{FN}\,*}$, the penalty for negatives $\delta^{\text{FP}\,*}$, and the weight for $\ell_1$ regularization term $C$. We fixed $\delta^{\text{FP}\,*} = 0.5$ because the balance between $\delta^{\text{FN}\,*}$ and $\delta^{\text{FP}\,*}$ is important. We performed the grid search on $\delta^{\text{FN}\,*} \in \{0.001, 0.005, 0.05, 0.5\}$ and $C \in \{10^n \mid n = -3, \ldots, -6\}$ for the whole dataset. Then, we chose $\delta^{\text{FN}\,*} = 0.05$ and $C = 10^{-5}$ at which good accuracy can be achieved. Tables S1 and S2 show the accuracy under varying $\delta^{\text{FN}\,*}$ and $C$ respectively, indicating that $\delta^{\text{FN}\,*}$ and $C$ are not strongly sensitive to the prediction accuracy.

### Supplementary Table

Table S1: Accuracy under varying $\delta^{FP*}$ with fixing $\delta^{FN*} = 0.5$ and $C = 10^{-5}$.

| | Contacts | | | Binding residues | | | Binding bases | | |
|---|---|---|---|---|---|---|---|---|---|
| $\delta^{FP*}$ | PPV | SEN | F | PPV | SEN | F | PPV | SEN | F |
| 0.001 | 0.550 | 0.505 | 0.513 | 0.686 | 0.684 | 0.670 | 0.698 | 0.631 | 0.647 |
| 0.005 | 0.555 | 0.504 | 0.514 | 0.698 | 0.684 | 0.674 | 0.698 | 0.630 | 0.645 |
| 0.05 | 0.539 | 0.525 | 0.519 | 0.664 | 0.688 | 0.657 | 0.684 | 0.655 | 0.652 |
| 0.5 | 0.432 | 0.571 | 0.476 | 0.509 | 0.652 | 0.554 | 0.529 | 0.656 | 0.570 |

Table S2: Accuracy under varying $C$ with fixing $\delta^{FP*} = 0.005$ and $\delta^{FN*} = 0.5$.

| | Contacts | | | Binding residues | | | Binding bases | | |
|---|---|---|---|---|---|---|---|---|---|
| $C$ | PPV | SEN | F | PPV | SEN | F | PPV | SEN | F |
| $10^{-6}$ | 0.541 | 0.519 | 0.517 | 0.669 | 0.677 | 0.656 | 0.674 | 0.646 | 0.646 |
| $10^{-5}$ | 0.539 | 0.525 | 0.519 | 0.664 | 0.688 | 0.657 | 0.684 | 0.655 | 0.652 |
| $10^{-4}$ | 0.504 | 0.531 | 0.501 | 0.622 | 0.686 | 0.631 | 0.661 | 0.663 | 0.644 |
| $10^{-3}$ | 0.265 | 0.392 | 0.293 | 0.340 | 0.509 | 0.379 | 0.362 | 0.498 | 0.391 |

Table S3: Structural profiles coding

| Residue | $\alpha$-helix | 3-helix | 5-helix | folded | $\beta$-turn | corner | curl | loop |
|---|---|---|---|---|---|---|---|---|
| | H | G | I | E | B | T | S | – |
| Base | External | Hairpin | Internal | Bulge | Multibranch | Stack | | |
| | E | H | I | B | M | S | | |

Table S4: Coding of simplified alphabets (10 groups)

| amino acids | LVIM | C | A | G | ST | P | FYW | EDNQ | KR | H |
|---|---|---|---|---|---|---|---|---|---|---|
| coding | L | C | A | G | S | P | F | E | K | H |

Table S5: Coding of simplified alphabets (4 groups)

| amino acids | LVIMC | AGSTP | FYW | EDNQKRH |
|---|---|---|---|---|
| coding | L | A | F | E |
